# Does the syrinx, a peripheral structure, constrain effects of sex steroids on behavioral sex reversal in adult canaries?

**DOI:** 10.1101/2023.04.19.537462

**Authors:** Ednei B. dos Santos, David M. Logue, Gregory F. Ball, Charlotte A. Cornil, Jacques Balthazart

## Abstract

We previously confirmed that effects of testosterone (T) on singing activity and on the volume of brain song control nuclei are sexually differentiated in adult canaries: females are limited in their ability to respond to T as males do. Here we expand on these results by focusing on sex differences in the production and performance of trills, i.e., rapid repetitions of song elements. We analyzed more than 42,000 trills recorded over a period of 6 weeks from 3 groups of castrated males and 3 groups of photoregressed females that received Silastic™ implants filled with T, T plus estradiol or left empty as control. Effects of T on the number of trills, trill duration and percent of time spent trilling were all stronger in males than females. Irrespective of endocrine treatment, trill performance assessed by vocal deviations from the trill rate versus trill bandwidth trade-off was also higher in males than in females. Finally, inter-individual differences in syrinx mass were positively correlated with trill production in males but not in females. Given that T increases syrinx mass and syrinx fiber diameter in males but not in females, these data indicate that sex differences in trilling behavior are related to sex differences in syrinx mass and syrinx muscle fiber diameter that cannot be fully reversed by sex steroids in adulthood. Sexual differentiation of behavior thus reflects organization not only of the brain but also of peripheral structures.

## Introduction

Sexual reproduction is often preceded by elaborate courtship displays of males that are used by females to assess the quality of potential mates. Bird song is a textbook example of such displays (Collins, 2004). Several song features that are thought to be reliable signals of male quality in songbirds have been identified in various species. These features include song repertoire size (Catchpole and Slater, 2018), stereotypy (Sakata and Vehrencamp, 2012), and the occurrence of ‘special’ syllables (Rehsteiner et al., 1998). By mating with males that produce songs displaying these characteristics, females can potentially obtain greater direct benefits, such as better parental care and access to larger territories with more resources, or indirect genetic benefits (better genes) for their offspring (Andersson, 1994; Kirkpatrick and Ryan, 1991).

Vocal features that are physically challenging to produce can also indicate male quality and these features have often been grouped under the term of vocal performance (Goller, 2022; Podos et al., 2009). Vocal performance has traditionally been measured as the rate of vocal production (number of vocalizations per unit time) but it can also be assessed by vocal consistency (the degree of variation across renditions of the same song elements) and by the fine-scale structure of trills (rapid repetitions of song elements in sequence). Podos proposed that physiological constraints result in a trade-off between trill rate and trill frequency bandwidth in the song of over 30 species of sparrows (Emberizidae) (Podos, 1997). Similar trade-offs have since been reported in several other species (reviewed in (Goller, 2022; Podos et al., 2009; Wilson et al., 1997). These tradeoffs emerge because there is a limit to the speed with which a bird can modulate the fundamental frequency of its song. Both trill rate and frequency bandwidth rely on frequency modulation over time. Therefore, as a bird approaches the limit of frequency modulation speed, any increase in trill rate must be offset by a decrease in frequency bandwidth, and *vice versa*. When plotted in an acoustic space defining trill bandwidth as a function of trill rate, the distribution of trills forms a right triangle with the hypotenuse defining a putative performance limit. Trills that are orthogonally closer to this optimal limit have lower vocal deviation (i.e., higher performance) and are more challenging to sing because they require faster movements or better coordination of the respiratory system, syringeal muscles and the vocal tract.

In canaries (*Serinus canaria*), females exhibit copulation solicitation displays more often and deposit more testosterone in eggs when exposed to playbacks of male songs containing trills composed of broad frequency bandwidth two-note syllables repeated at a fast rate of at least 15 elements s^-1^ that were labeled ‘sexy’ or ‘A’ syllables (Vallet et al., 1998; Vallet and Kreutzer, 1995). Suthers and colleagues investigated the patterns of airflow in the syrinx (the avian vocal organ) during song production and demonstrated that canaries employ two different respiratory mechanisms to produce trills (Suthers et al., 2012). The rapid production of syllables is normally constrained by the respiratory system that limits the rate at which air can be replaced. Canaries circumvent this limitation by taking ‘mini-breaths’ between syllables, allowing them to sing trills of protracted duration. However, to produce faster trills canaries shift to ‘pulsatile respiration’, in which expiration is sustained for the entire duration of trills, thus limiting trill duration to the available air supply. Thus, long sexy trills are both preferred by females and challenging for males to produce.

The two independent sound sources in the syrinx associated with the left and right bronchi are controlled by a minimum of 4 pairs of muscles that are among the fastest known muscles in vertebrates (Elemans et al., 2008). These muscles are partially controlled by steroid hormones. Both androgen receptors and estrogen receptor β are expressed in the syringeal muscles of male and female zebra finches (*Taeniopygia gutatta*) (Veney and Wade, 2005). Furthermore, a study in male canaries in which syringeal androgen receptors were blocked with bicalutamide (an androgen receptor antagonist that does not cross the blood-brain barrier) reported a significant decrease in syrinx mass associated with a reduction in trill performance and complexity (Alward et al., 2016). Syrinx muscle fibers are additionally controlled by the singing activity and muscles training (Adam and Elemans, 2019). Sex differences in syrinx mass and muscle fibers composition have also been reported in several songbird species (Christensen et al., 2017; Prince et al., 2011; Wade and Buhlman, 2000). However, studies directly relating sex differences in syrinx mass or anatomy to sex differences in singing behavior are scarce.

Canaries have frequently been used as a model to study effects of sex steroids on singing behavior (see reviews in (Ball and Balthazart, 2007; Schlinger and Brenowitz, 2017)). In a previous study we compared the role of testosterone (T) on singing activity and brain song control nuclei anatomy in both males and females (Dos Santos et al., 2022). That study confirmed that the sexes respond differentially to T and there is a limit in the capacity of females to respond to T in the same manner as males. Here we expand on these results by focusing on sex differences in the production and performance of trills. First, we analyze in detail the effects of T on trill production and trill features. Then we assess sex differences in T effects on performance measured by vocal deviations from the trill rate versus trill bandwidth trade-off. Finally, we explore how sex and inter-individual differences in syrinx mass and syringeal muscle fiber diameter relate to trill acoustic features and performance. Although a number of studies have demonstrated that expression of a given behavior can be limited by peripheral structures, since the discovery of sex differences in brain structure and function in songbirds (Nottebohm and Arnold, 1976) and then in many other species (Tobet and Fox, 1992), when trying to explain sex differences in behavior, it is usually at the brain level that answers are searched for. Together the present data suggest that sex differences in trilling behavior are caused, at least in part, by sex differences in syrinx mass and muscle fiber diameter that cannot be reversed by sex steroids in adulthood.

## Material and methods

One-year old Fife Fancy canaries (24 females and 24 males) were acquired from a local commercial breeder in Belgium and sexed by PCR at the Behavioral Ecology and Ecophysiology lab of the University of Antwerp, Belgium (Griffiths et al., 1998). All birds were housed in groups of six in visually (but not acoustically) isolated cages under an 8 L: 16 D photoperiod at the animal facility of University of Liege, Belgium, with food, water, bath and grit ad libitum during the entire experiment. They were also fed egg food twice per week.

All experimental procedures complied with Belgian laws concerning the Protection and Welfare of Animals and the Protection of Experimental Animals, and experimental protocols were approved by the Ethics Committee for the Use of Animals at the University of Liege (Protocol number 2027).

### General procedure

Six weeks after their arrival in the laboratory, all males were castrated and females were laparotomized under general isoflurane anesthesia to confirm photoregression of their ovary as described before (Dos Santos et al., 2022; Goldman and Nottebohm, 1983; Hartog et al., 2009; Madison et al., 2015; Shevchouk et al., 2017). Adult female canaries with regressed gonads are commonly used to investigate the effects of T on song system neuroplasticity (Hartog et al., 2009; Louissaint et al., 2002; Yamamura et al., 2011). Three weeks later, all birds were weighed and the size of their cloacal protrusion area, an androgen-dependent structure (Alward et al., 2013; Appeltants et al., 2003; Tramontin et al., 2003) was measured with calipers before they received subcutaneous implants made of Silastic™ tube (Dow corning, Midland, MI, USA; Degania Silicone; internal diameter 0.76 mm, external diameter 1.65 mm, length 12 mm) filled over a 10 mm length with either crystalline T or estradiol (E2) or left empty as a control (C). These implants maintain concentrations of T and E2 that are in the high physiological range of behaviorally active of doses (Appeltants et al., 2003; Cornez et al., 2020; Leboucher et al., 1994; Madison et al., 2015; Sartor et al., 2005).

Birds from each sex were randomly assigned to three experimental groups that received either two empty implants (C group), one T and one empty implant (T group) or one T and one E2 implant (T+E2 group). Birds were then moved into 8 cages (4 with males and 4 with females), each housing two C, two T and two T+E2 birds

We included a group treated with E2 in addition of T because previous studies have reported that aromatase activity and that the induction of this enzymatic activity by T is lower in the female than in the male brain and this consequently might explain the limited response of females to exogenous T (for more detail, see (Dos Santos et al., 2022)).

### Song recording

Once every week for 6 weeks, birds were moved overnight individually inside custom-built sound-attenuated boxes. Vocalizations produced by each bird were recorded for 3 h starting immediately after lights on (0900 h) on the next morning using custom-made microphones (Projects Unlimited/Audio Products Division) and an Allen & Heath ICE-16 multichannel recorder. The procedure was repeated once a week for a total of 6 weeks starting one week after steroid implantation.

Sound files were acquired and saved as a .wav file by Raven v1.4 software (Bioacoustics Research Program 2011; Raven Pro: Interactive Sound Analysis Software, Version 1.4, Ithaca, NY: The Cornell Lab of Ornithology) at a sampling frequency of 44,100 Hz. Sound files were then analyzed with a MATLAB script developed for canary song analysis by Ed Smith and Robert Dooling (Department of Psychology, University of Maryland at College Park, MD). Songs were defined as vocalizations at least 30 dB above background noise that were at least 1 s long, and were preceded and followed by at least 0.4 s of silence. The present report is based on these recordings (24 male and 24 female Fife fancy canaries). Recordings were already used to analyze the sexually differentiated effects of sex steroids on singing behavior and brain plasticity. These results have been published separately (Dos Santos et al., 2022).

### Tissue collection

Six weeks after the beginning of the steroid treatment, birds were weighed, their cloacal protrusion area was measured again and they were deeply anaesthetized before being euthanized by decapitation. Their brain was dissected out of the skull and fixed with 5% acrolein, cryoprotected in sucrose, frozen on dry ice and stored at -80°C (See (Dos Santos et al., 2022) for more detail). The syrinx of each subject was also extracted, fixed with acrolein, frozen on dry ice and stored at -80°C until used.

### Tissue processing

All syringes were defrosted and dissected by one cut of the trachea just dorsal to the syringeal muscles and two cuts ventral to the third cartilaginous ring of the bronchi. Each syrinx was then weighed to the nearest milligram. They were then cut with the cranial aspect facing up on a cryostat (Thermo ScientificTM CryoStarTM NX70) in 15 µM coronal sections that were mounted on superfrost microscopic slides. Sections were stained with hematoxylin-eosin and coverslipped with permount. The section containing the largest cross-sectional area was photographed under a microscope at a 10X objective for each bird. A square (200×200 µM) was overlaid in each of the 4 quadrants of the syrinx photomicrograph (quadrants 1 and 2, left and right ventral side, including the muscles tracheobronchealis ventralis and syringealis ventralis ; quadrants 3 and 4, left and right dorsal side, including the muscles tracheobronchealis dorsalis and syringealis dorsalis) using the FIJI version of ImageJ software (Schindelin et al., 2012). All fibers located within the square were measured (n= 3,401 fibers for all birds). The widest and narrowest diameter of each fiber was measured. These 2 measures were averaged and considered as the mean diameter which was then averaged across all quadrants and each bird before statistical analyses. The distribution of muscle diameters within each experimental group was also investigated.

### Trill analysis

Trills were identified within songs and defined as sequences of similar elements repeated at least 4 times and separated by silence intervals of no more than 0.003 s. with the help of a MATLAB routine especially developed by Ed. Smith and Rober Dooling (Department of Psychology, University of Maryland at College Park) for the analysis of canary song. The script computed the following metrics: total number of trills, percent time spent trilling (within songs), trill duration, number of segments (per trill), segment duration, number of fast trills (trills with a rate of at least 17 syllables s^-1^), interval duration (between segments), spectral distance (between segments), trill entropy, trill bandwidth, trill center frequency and trill power. One female bird died of natural causes during the experiment and was recorded only for a 5-week period.

### Performance analysis

Because the production of trills might be constrained by a number of physical factors, we also quantified the trill rate (TR) vs. mean trill bandwidth (TBW) trade-offs, while controlling for variation attributable to individuals. Trill rate (TR) was calculated as the number of segments minus one divided by the time from the beginning of the first segment to the beginning of the last segment. The final segment was excluded from this measure because it is impossible to define the duration of the silent gap that follows this last segment and this would bias estimates of trill rate for song with fewer segments. Mixed quantile regressions (99^th^) were used to test for acoustic trade-offs (Logue et al., 2020; Wilson et al., 1997). A quantile regression analysis generates a linear function to estimate a defined quantile variable Y over a range of X (Cade and Noon, 2003). All models were run with both random intercepts and random slopes, and to account for the non-independence of multiple data points from the same individuals, we used individual id as a random variable. We then calculated deviation scores (DS) as the orthogonal distance from the quantile regression line (Podos, 2001) defined by: DS=(TWB intercept + TBW slope X TR –TBW)/sqrt (1+TBWslope^2^) X -1

### Statistical analysis

Unless otherwise mentioned, data were analyzed by one-or two-way analyses of variance (ANOVA) or by two-way mixed model analysis if a few data points were missing with the three experimental groups, as appropriate. Post hoc comparisons were performed with the Tukey’s multiple comparisons test. Nominal data were analysed by the χ2 and Fisher exact probability tests. Statistical analyses were performed using R Studio (Team, 2021) and GraphPad Prism version 8.4 for Mac (GraphPad Software, San Diego, California USA). We used the lqmm package (Geraci, 2014) to calculate mixed quantile regressions, and the package ggplot2 (Wickham and Chang, 2008) to graph the trill distributions with semi-transparent points and to fit the mixed quantile regression line. Effect sizes (partial eta square η_p_^2^) are represented by the ratios of the relevant sums of squares in the two way ANOVA. We used an alpha level of .05 for all statistical tests. All data are represented here by their mean ± SEM and when feasible, individual data points are also presented.

## Results

Additional results are available in online Supplementary Results.

### Analysis of trills

A total 864 hours of recordings (3h per week for 6 weeks for 48 experimental subjects) were analyzed by the MATLAB script that detected and quantitatively characterized a total of 42,198 trills. Twelve dependent variables were analyzed by three types of two-way ANOVAs. Data for each dependent variable were first averaged for each subject across the 6 weeks of experiment and analyzed by a two-way ANOVA with endocrine treatment (3 conditions: C, T and T+E2) and sex (male or female) as independent factors (Fig. 1, Fig. S1 and Table S1).

**Fig. 1.**
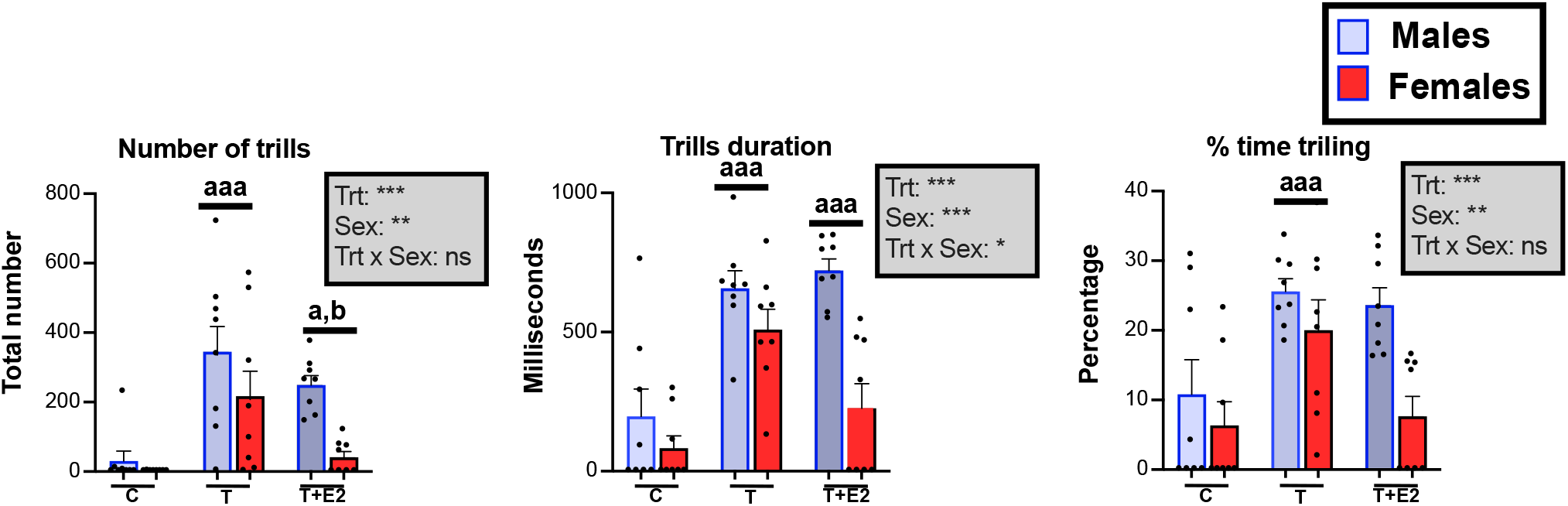
Number of trills (A), trill duration (B) and percentage of time spent trilling (C) in male and female canaries that were treated with Silastic™ implants filled with testosterone (T) or with testosterone plus estradiol (T+E2) or left empty as control (C). Bar graphs represent the mean ± SEM of individual results that are the average of data collected during the 6 weeks of recording. Individual data points are also indicated. Data were analyzed by two-way ANOVA with treatment (Trt) and Sex of the subjects as independent factors and results are summarized in the insert for each panel. (***=p<0.001, **=p<0.01, *=p<0.05, ns= not significant). Significant effects of treatment were further analyzed by Tukey’s multiple comparison tests and their results are expressed as follows: aaa= p<0.001 versus C group, a (or b)= p<0.05 versus C (or T) group.

Over the entire experiment, steroid treatments significantly affected many aspects of the trills including their total number, percent time trilling, trill duration (Fig. 1) and also the number of segments per trill, the segment duration and the number of fast trills containing more than 17 elements per second (Fig. S1; see Table S1 for detail of statistical analyses). In most cases, p*osthoc* analyses identified significant differences between the C and the T and/or the T+E2 groups (see detail in Fig. 1 and Fig. S1). These analyses also indicated a significantly lower total number of trills in the T+E2 groups compared to the T groups.

The ANOVAs also detected significant sex differences for three variables: the total number of trills, percent time trilling, and trill duration (Fig. 1, Table S1). A significant interaction between treatment and sex was observed for trill duration only (sex difference was larger in the T+E2 condition than in the two other conditions).

Initially, females and castrated males did not produce trills; trills appeared progressively during the treatment with steroids (Fig S2). Separate two-way ANOVAs within each sex with time and treatment as factors confirmed the presence of treatment and time effects for several trill features in males (Fig. S2 and Table S2). Furthermore, two-way ANOVAs within each treatment with time (weeks) and sex as factors) confirmed the presence of a sex difference in trill numbers, percent time trilling and trill duration in the T+E2 birds (Fig. S2 and Table S3).

### Trill rate (TR) versus Trill Bandwidth (TBW) trade-off and performance analysis

The analysis of the 42,198 trills produced by all experimental birds across the 6 weeks of experiment identified a trade-off between TR and mean TBW, presumably reflecting physiological limits on frequency modulation and respiration in canary trill production (Fig. 2). A quantile regression analysis on the pooled data from the 35 subjects that produced trills during the experiment indicated the presence of statistically significant, negatively sloping upper boundaries for TR vs. TBW (intercept = 2.98, slope = −1.6, pslope < 0.001, Fig. 2*A*).

**Fig. 2.**
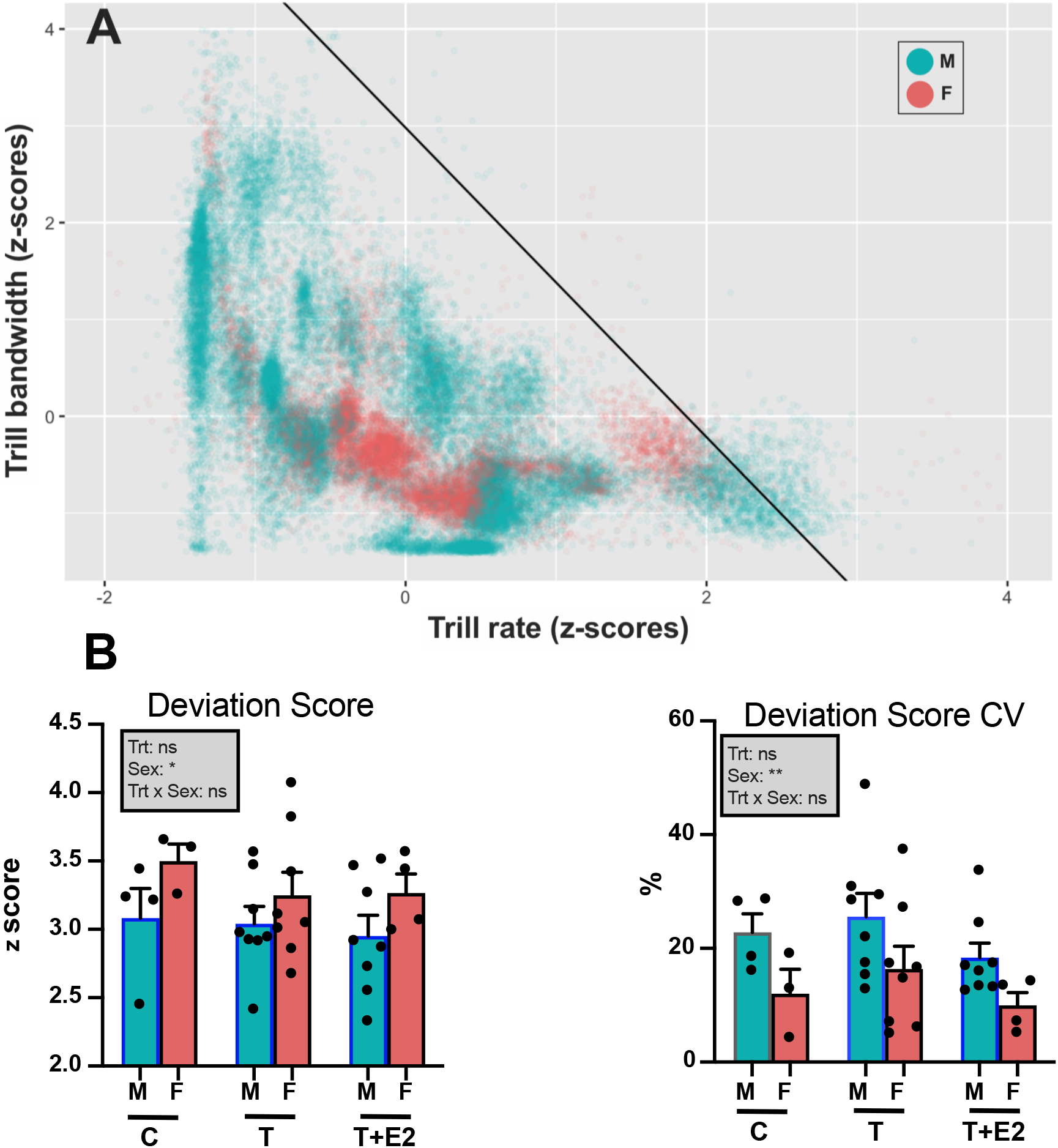
Trade-off between trill rate and trill bandwidth in songs of male (M) and female (F) canaries that were treated with Silastic implants filled with testosterone (T), with testosterone plus estradiol (T+E2) or left empty as control (C). Panel (A) shows the position of each of the individual trills with respect to the quantile regression line with male data points indicated in turquoise and female points in red. Panel (B) shows the mean ± SEM deviation scores and coefficient of variation of these scores for each bird in the six experimental groups. Results of the two-way ANOVA of these data are summarized in the insert (*=p<0,05, **=p<0,01, ns= non-significant).

The deviation score (orthogonal distance from the regression line) for each individual trill was then calculated, averaged within each subject and these individual scores were then compared by a two-way mixed model ANOVA with the sex and treatment of the birds as independent factors. This analysis identified a significant sex difference in the deviation scores (Fig. 2*B*), with males on average having a smaller orthogonal distance from the quantile regression line (F_1,29_= 4.487, p=0.043, η_p_^2^=0.21). There was however no effect of the endocrine treatments (F_2,29_= 0.460, p=0.636, η_p_^2^=0.03) and no sex by treatment interaction (F_2,29_= 0.174, p=0.841, η_p_^2^=0.01) related to these scores.

In addition, deviation scores were more variable in males than in females reflecting the fact that male trills were more broadly distributed across the entire acoustic space. Analysis of the deviation score coefficients of variation accordingly revealed an overall sex difference (F_1,29_= 7.778, p=0.009, η_p_^2^=0.21), but again no significant effect of treatment (F_2,29_= 1.750, p=0.192, η_p_^2^=0.11) and no interaction between sex and treatments (F_2,29_= 0.034, p= 0.966, η_p_^2^=0.00).

One interesting feature of data illustrated in Fig. 2*A* is that the points representing individual trills were not randomly distributed in space but rather were grouped in relatively discrete clusters. Additional plots of these trills in the TR vs. TBW space indicated that these clusters are different for males and females (Fig. S3) as well as in individual subjects (Fig. S4), suggesting they relate to different types of syllables used in the trills. This hypothesis was confirmed by the qualitative analysis of sonograms presented in Figure 3.

**Fig. 3.**
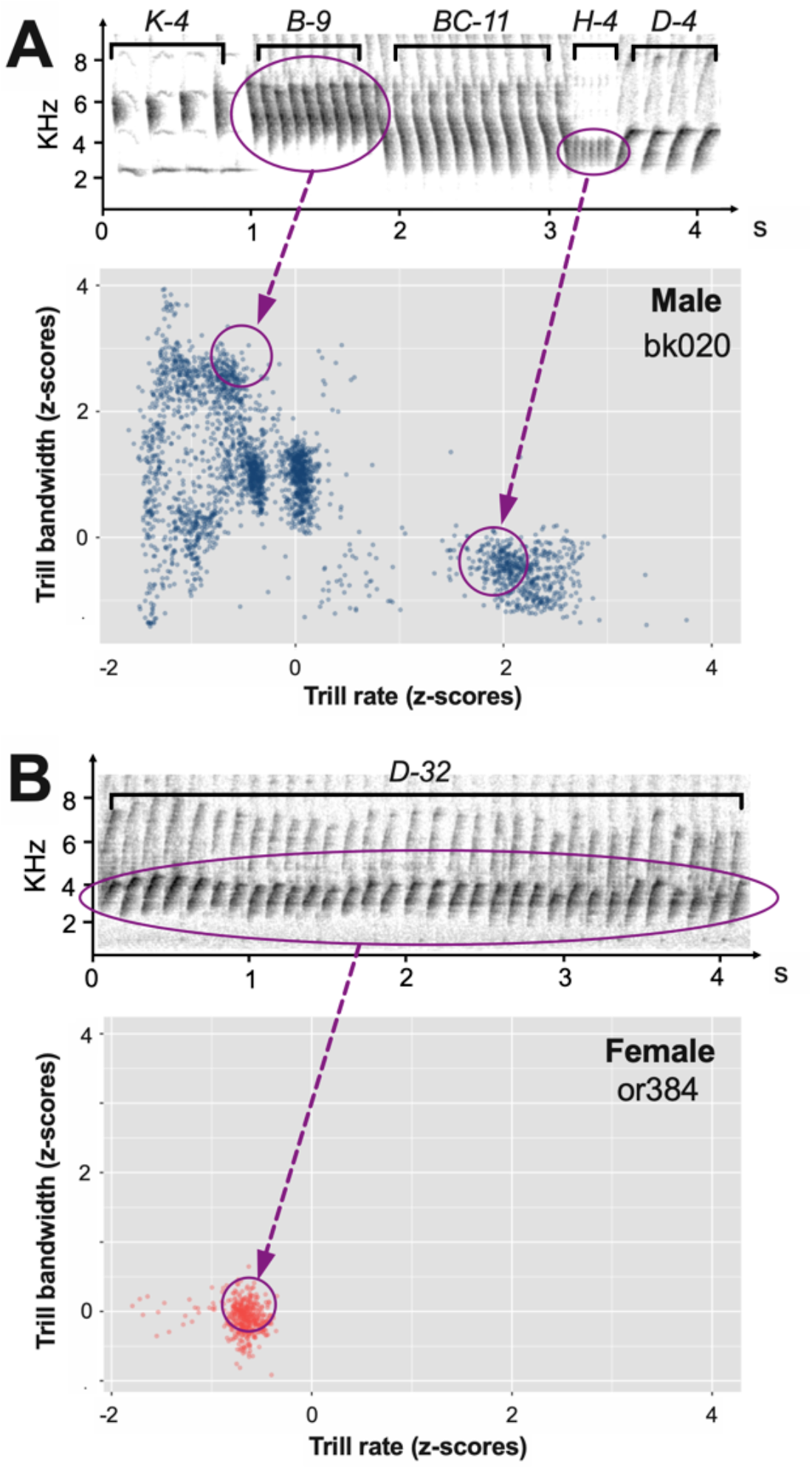
Example of sonograms from a typical song in a male (A) and a female (B) canary illustrating the repetition of a same element within long trills in females and use of various elements in a male. In the trade-off plots, female or384 displays a single cluster of data points while male bk020 shows multiple clusters corresponding to the different elements of the trills. Some of these elements are easily identified by the combina-tion of their bandwidth and rate (circles and arrows).

### Effects on syrinx mass and structure

In a previous study, we showed that T or T+E2 increase the volume of three song control nuclei (HVC, RA and Area X) in both sexes but values remained significantly smaller in females than in males (Dos Santos et al., 2022). In parallel, T increased syrinx mass in males but not in females (Dos Santos et al., 2022).

Correlatively, the syrinx muscles fiber diameter was differentially increased by T in males and females (Fig. 4D) (Effect of treatment: F_2,32_= 3.87, p<0.001, η_p_^2^0.069; effect of sex F_1,32_= 58.05, p<0.001, η_p_^2^=0.64; interaction: F_2,32_= 16.40, p<0.001, η_p_^2^0.51). While syrinx fibers average diameter was similar in control males and females (Post hoc Sidak’s multiple comparisons: p=0.999), this diameter was significantly larger in males than in females in the two T-treated groups (Sidak test: p<0.001 in both cases Fig. 4D). A similar differential increase in males and females of the syrinx fiber diameter was actually observed for each of the 4 quadrants of the syrinx (see Fig. S5A). Correlatively this increased fiber size resulted in a decrease in the density of fibers (numbers per unit surface) in the two male groups treated with T (Effect of treatment: F_2,32_= 7.77, p=0.002, η_p_^2^=0.33; effect of sex F_1,32_= 6.48, p=0.02, η_p_^2^=0.17; interaction: F_2,32_= 2.61, p=0.089, η_p_^2^=0.14; Fig. S5B).

**Fig. 4.**
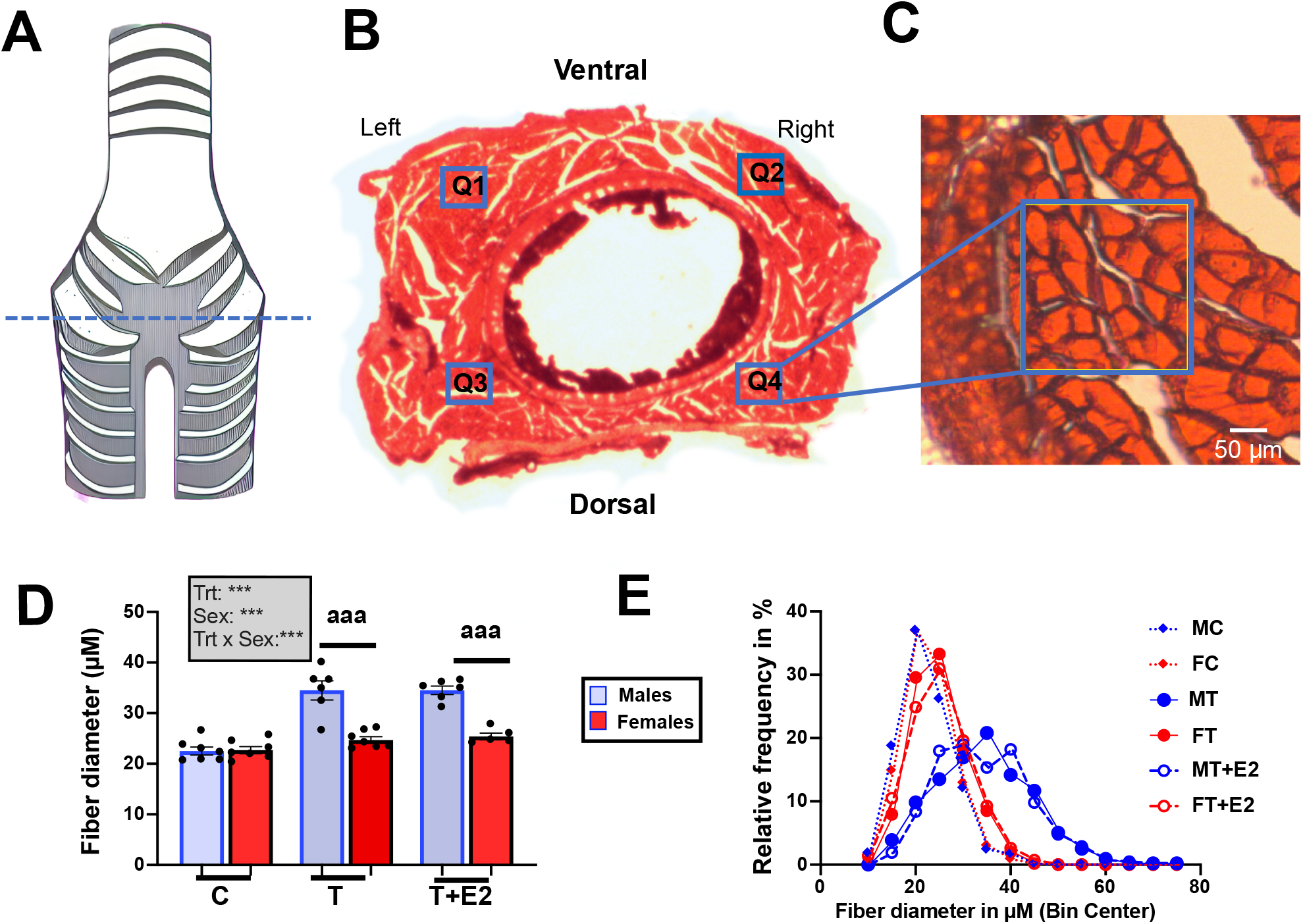
Effects of exogenous hormones on syrinx muscle fiber diameter and density of fibers per unit surface. **(A)** Schematic view of the syrinx with the plane of section indicated by the dotted line, **(B)** Histological section through the syrinx showing the 4 quadrants(Q1-4), **(C)** Example of fibers viewed at higher magnification, **(D)** Mean diameter of the syrinx muscle fibers across the 4 quadrants, **(E)** Overall distribution of fiber sizes measured in the 4 quadrants in the 6 experimental groups. Data in panel D were analyzed by two-way ANOVA and results are summarized in the insert (***=p<0.001, **=p<0.01, *=p<0.05, ns= non-significant).

Plotting the distribution of all fibers measured in the 4 quadrants in each group separately revealed a shift of the distribution to the right (towards larger diameters) for the two groups of males that had been treated with T (T and T+E2; Fig. 4E) while this effect was not present in females; their distribution still overlapped with the distribution in control birds, never or very rarely reaching a size over 45 µM. This presence of larger fibers concerned all males in the T groups: fibers with a mean diameter larger than 45 µM were observed in 6 males out of 6 in the T and T+E2 groups. In contrast these larger fibers were completely absent in control birds (0 out of 7 in MC and FC) and rare in females treated with T (1/7 in females T and 2/5 in females T+E2). Overall, this distribution was significantly different from random (χ2 test= 29.39, df=5, p<0.001). Analyses confined to each treatment separately indicated the presence of a significant difference between T males and T females (6/6 vs. 1/T ; Fisher exact probability test : p=0.005) but not in T+E2 birds (6/6 vs 2/5, p=0.061) nor in controls (0/T vs. 0/7 ; p>0.999).

The number of large fibers (>45 µM) detected in the quantification square (200 × 200 µM) was similarly affected by the sex of the birds (F_1,32_=49.25, p<0.001, η_p_^2^=0.61), the endocrine treatments (F_2,32_=14.56, p<0.001, η_p_^2^=0.48) and the interaction between the 2 factors (F_2,32_=13.16, p<0.001, η_p_^2^=0.45). Posthoc Tukey tests within each sex demonstrated a significant increase in the two males groups treated with T by comparison with the controls (C :0±0, T :10.17±2,65, T+E2 : 9.83±1.19, p<0.001 in both cases) while such an effect was not observed in females (C :0±0, T :0.14±0.14, T+E2 : 0.40±10.24, p<>0.968 in both cases).

### Correlations of trilling behavior with syrinx mass and structure

To further investigate whether these morphological sex differences could explain the steroid resistant sex differences in trilling behavior, we next examined correlations between Syrinx muscle diameters and trilling behavior in males and females separately. This analysis revealed in males significant correlations between numbers of trills, trill duration and percentage of time trilling, on the one hand and syrinx muscles diameters, on another hand. These correlations were not detected females with the exception with a low correlation between numbers of trills and fiber diameters (Fig. 5).

**Fig. 5.**
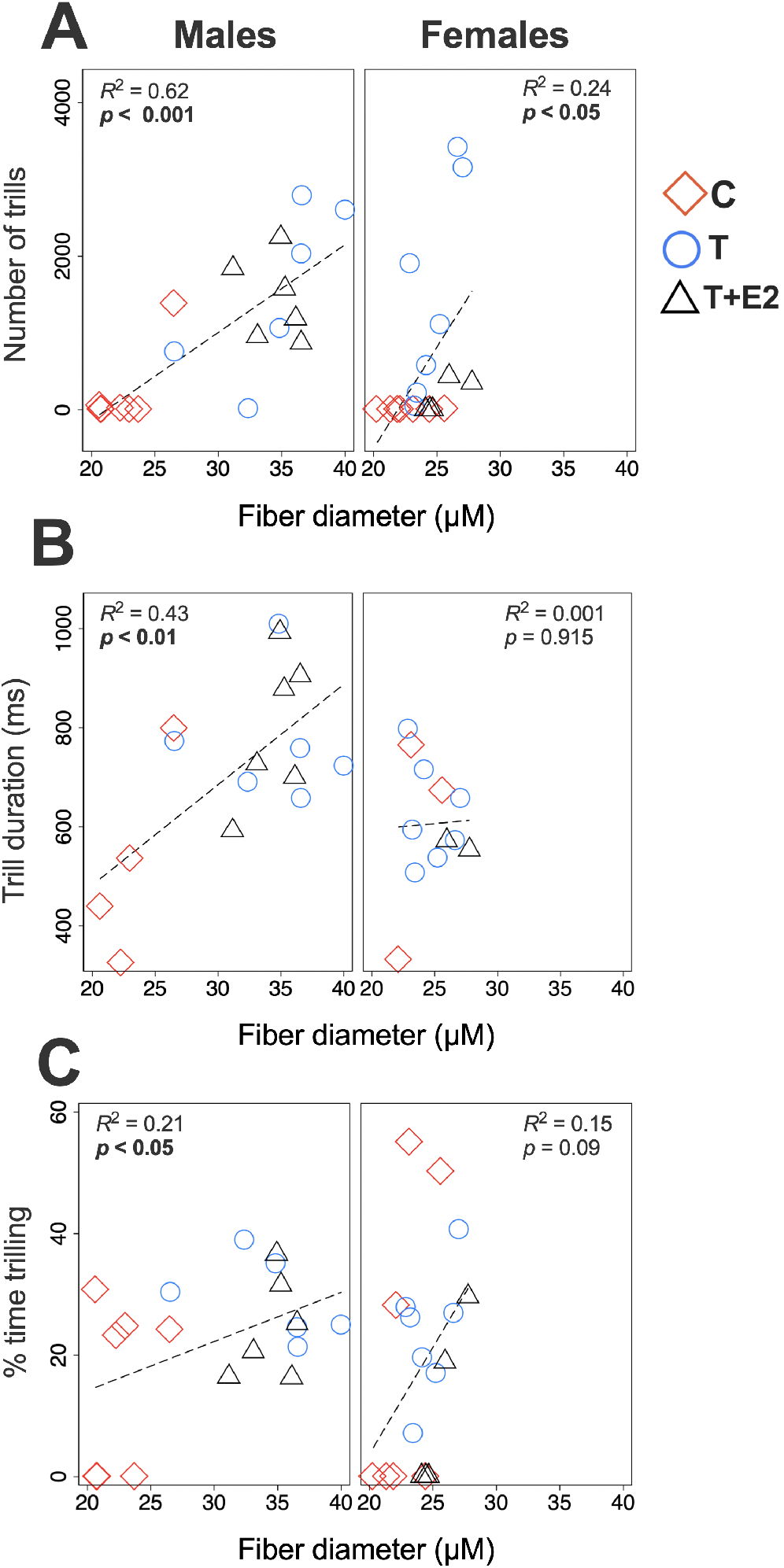
Correlations between trilling behavior with syrinx muscle diameters in males and females.

Then we developed multiple regression models for males and females in which the three measures of trilling were the dependent variable and muscle fibers diameter (Table 1) was the independent variables (predictors) together with the whole body mass.

**Table 1.** Multiple linear regression models of trilling behavior (rate, duration, percent time trilling) on predictor variables (fiber diameter with whole body mass), including for each predictor the estimate and size (η^2^)*.

|  | Males |  |  |  | Females |  |  |  |
| --- | --- | --- | --- | --- | --- | --- | --- | --- |
| **Number of trills** | *(F2,16 = 13.52, Adj. R2 = 0.58, p < 0.001)\*\*\** |  |  |  | *(F2,16 = 3.13, Adj. R2 = 0.19, p = 0.071)* |  |  |  |
| **Predictors** | **Estimate (SE)** | **t** | **p** | **$\eta^2$** | **Estimate (SE)** | **t** | **p** | **$\eta^2$** |
| Fiber diameters | 112.4 (25.8) | 4.35 | <0.001 *** | 0.63 | 300.5 (120.3) | 2.50 | 0.024 | 0.28 |
| Body mass | 11.3 (80.5) | 0.14 | 0.89 | 0.001 | 124.5 (132.8) | 0.94 | 0.363 | 0.05 |
| Intercept | -2586.2 (1359.7) | -1.9 | 0.075 | ---- | -9115.8 (4502) | -2.02 | 0.060 | --- |
| **Trill duration** | *(F2,13 = 7.73, Adj. R2 = 0.47, p = 0.006)\*\** |  |  |  | *(F2,9 = 0.33, Adj. R2 = 0.06, p = 0.725)* |  |  |  |
| **Predictors** | **Estimate (SE)** | **t** | **p** | **$\eta^2$** | **Estimate (SE)** | **t** | **p** | **$\eta^2$** |
| Fiber diameters | 25.9 (6.6) | 3.93 | 0.002 ** | 0.54 | 1.5 (21.9) | 0.07 | 0.946 | 0.0005 |
| Body mass | -32.3 (17.7) | -1.83 | 0.091 | 0.2 | -20.5 (25.4) | -0.81 | 0.439 | 0.068 |
| Intercept | 540.5 (315.5) | 1.71 | 0.11 | ---- | 965.2 (750.4) | 1.29 | 0.23 | --- |
| **% time trilling** | *(F2,16 = 2.25, Adj. R2 = 0.12, p = 0.137)* |  |  |  | *(F2,16 = 1.47, Adj. R2 = 0.05, p = 0.25)* |  |  |  |
| **Predictors** | **Estimate (SE)** | **t** | **p** | **$\eta^2$** | **Estimate (SE)** | **t** | **p** | **$\eta^2$** |
| Fiber diameters | 0.9 (0.5) | 2 | 0.063 | 0.22 | 3.7 (2.2) | 1.68 | 0.112 | 0.15 |
| Body mass | -0.6 (1.4) | -0.44 | 0.665 | 0.01 | 0.6 (2.4) | 0.24 | 0.811 | 0.004 |
| Intercept | 7.2 (24.1) | 0.3 | 0.768 | ---- | -82.4 (81.1) | -1 | 0.33 | --- |

\* Effect size interpretation for $\eta^2$ : small=0.01; medium=0.06; large=0.14.

The independent variables accounted for more than 50% of the variance in trills and trill duration in males but less than 28% in females (Table 1). There was a significant positive relationship between the number of trills and fiber diameters (*t* = 4.35 *p* < 0.001) and between trill duration and fiber diameters (*t* = 3.93, *p* = 0.002). In females, these relationships were not significant (Table 1). However, we did not find a significant relationship between the percent time trilling and fiber diameters in either sex even if this effect was close to significance imales.

## Discussion

The present study shows that treatments with sex steroids (T or T+E2) markedly increase the number and several features of trills produced by both male and female canaries. T also increases the syrinx mass and syrinx muscles fiber diameter in males but not females and inter-individual differences in syrinx mass correlate positively with multiple aspects of trill production in males but not females. These results are in line with our previously reported data on the effects of steroid treatments on song production (Dos Santos et al., 2022) and additionally strongly suggest that the sex differences in trill production are caused, at least in part, by the sex difference in the peripheral vocal organ, the syrinx. The prominent sex differences in these effects deserve further discussion.

### Sex differences in trill production

Analysis of the entire data set over 6 weeks (Fig. 1, S1 and Table S1) revealed significant sex differences for three variables: the total number of trills, percent time trilling, and trill duration. A significant interaction between treatment and sex was also observed for trill duration (sex difference was larger in the T+E2 condition than in the two other conditions). Qualitative inspection of the data indicated that sex differences were mostly the result of larger effects of T and T+E2 in males than in females and accordingly, analysis of the effects of treatments week by week (Fig. S2 and Table S3) showed that sex differences are exclusively present in the T or T+E2 condition. Note also that trill production increased soon after the initiation of steroid treatments and plateaued after 3 or 4 weeks of treatment in both sexes. These results indicate that responses of females to T treatments are not slower than those of males, and also that the sex differences in responses are not due to the lower aromatization of T into estrogens in females since addition of exogenous E2 does not suppress the sex difference (Dos Santos et al., 2022).

Sex differences were particularly prominent for “fast trills” which contains more than 17 segments per second). Fast trills were almost completely absent in the T+E2 female vocalizations (Fig. S1). Female canaries exhibit copulation solicitation displays with higher frequency when exposed to playbacks of male songs containing ‘sexy’ syllables (Vallet et al., 1998; Vallet and Kreutzer, 1995). Additionally, the expression of immediate early genes *ZENK* and *c-Fos* in two auditory forebrain regions, the caudal mesopallium and nidocaudal mesopallium (areas analogous to secondary auditory cortices in mammals), is higher in female canaries exposed to male songs including ‘sexy’ trills than to songs without these sexy trills (Leitner et al., 2005; Monbureau et al., 2015), although these responses may depend on the acoustic context (Haakenson et al., 2019).

### Sex difference in trill performance

The TR versus mean TBW trade-off in canaries agrees with previous studies reporting similar findings for different species (see reviews in (Podos and Sung, 2020; Wilson et al., 1997). This trade-off probably results from constraints in frequency modulation and in rate of syllable repetition (Logue et al., 2020). The deviation scores from the TR vs. TBW trade-off were sexually differentiated (Fig. 2*B*), with males on average having a smaller but more variable orthogonal distance from the quantile regression line than females. However, there was no significant effect of the endocrine treatments and no sex by treatment interaction related to these scores. The higher dispersion in the trade-off space of trills produced by males compared to females (Fig. 2*A*) is especially prominent for the Y axis (trill bandwidth). The trill data points from females are mostly positioned in one or two clusters at a relatively low bandwidth.

This more variable trill bandwidth in males is functionally important since females have a preference for syllables with broad bandwidth (Draganoiu et al., 2002). They increase their copulation solicitation displays when exposed to syllables with a broad bandwidth broadcasted at an artificially increased rate. This preference for supranormal vocalizations in terms of bandwidth and repetition rate is consistent with the notion that these features are honest-signals of male quality (Draganoiu et al., 2002) that vary across male subjects while females lack the capacity to modulate trills to the same extent as males. This raises the obvious question of the mechanism(s) mediating this behavioral sex difference.

### Correlations with syrinx mass and structure

We previously reported that syrinx mass increases in response to steroid treatments in males but not in females (Dos Santos et al., 2022). These observations are indeed consistent with previous work that has shown that although testosterone does increase syrinx mass in adult songbirds, variation in T cannot explain the general sex differences affecting this structure and its function (reviewed in (Adam and Elemans, 2019). Here we show that syrinx mass and syringeal muscle fiber diameter are positively correlated with the number of trills and trill duration in males but not in females. Similarly, the diameter of syringeal muscle fibers increases in response to T in males but not in females. Larger diameters of syringeal muscles are known to be linked to the ability to produce faster rates of song (Christensen et al., 2017). These differences in the response to T in the male *vs*. female syrinx suggest that aspects of the syringeal structure that vary in males and females in response to T relate to sex differences in trilling rate and performance. Differences in syrinx morphology have already been proposed to explain sex differences in vocal behavior. In European Starlings (*Sturnus vulgaris*), for example, males have greater muscle mass in the syrinx, sing at higher rates and have larger vocal repertoires than females (Prince et al., 2011).

Sex differences in syrinx morphology might be explained by differences in testosterone sensitivity, mediated by androgen receptor (AR) expression in the syringeal muscles. In zebra finches, AR expression, as measured by *in situ* hybridization of the corresponding mRNA, is indeed denser in male than in female syringes (Veney and Wade, 2004, 2005). In songbirds, androgen sensitivity in specific parts of the song control system and periphery mediates the display of physically elaborate sexual displays (Fuxjager et al., 2015) and AR expression in the syrinx muscles is thus likely a sexually selected trait. Sexual selection might have increased the sensitivity of syringeal muscles to androgenic hormones specifically in males facilitating the production of vocal traits that are favored by females. This hypothesis is reinforced by a study in male canaries assessing effects of blocking androgen action in the syrinx by a treatment with the anti-androgen bicalutamide that does not cross the blood-brain barrier and thus cannot affect the brain but can act on peripheral structures such as the syrinx. Bicalutamide decreased the trill repetition rate without affecting other song features such as song rate, therefore linking androgen action in the syrinx directly to features of song that are known to be sexually selected (Alward et al., 2016). Additional studies can asses whether those birds that are able to produce trills at higher rates express greater levels of androgen receptors in the syrinx. Overall the significant correlation between trill production and syrinx mass or syrinx muscle fibers diameter in males supports the view that the syrinx limits trill production and explains the trilling sex difference.

An alternative, non-mutually exclusive, explanation for the sex difference in syrinx mass relates to the fact that T-treated males sang many more songs than T-treated females so that the larger syrinx mass in males could be activity-dependent, i.e., the result of more exercise (Alvarez-Borda and Nottebohm, 2002; Alward et al., 2013; Alward et al., 2016; Li et al., 2000; Maxwell et al., 2021). Indeed, experiments in Starlings have identified a 20–40% decrease in syringeal muscle mass after vocal denervation (Prince et al., 2011), suggesting a connection between vocal activity and syrinx mass.

The sex differences in trill production could obviously also result from differences in song control nuclei anatomy. Our previously published study based on the same subjects had indeed identified stable sex differences in three song control nuclei that were not suppressed by the treatments with T or T+E2 (Dos Santos et al., 2022). However, the present study reveals that multiple sex differences in trilling behavior and performance as well as individual differences among males are correlated with and might be caused, at least in part, by differences in syrinx mass and structure. The proposed relationship between a sophisticated learned vocal behavior and a peripheral vocal organ is reminiscent of the sex dimorphism in the vocal-fold and vocal tract lengths that emerges during adolescence and is mediated by testosterone in humans. In males, during adolescence, mean voice *F*_*0*_ and formant frequencies decrease around 50-60% and 80-90% respectively when compared to female values (Owren et al., 2007).

The modern field of the neuroendocrine basis of sexual differentiation was established by the seminal paper of Phoenix et al. (Phoenix et al., 1959) who argued that perinatal steroid hormones organize the brain in a male-typical or female-typical manner that in many cases can not be reversed in adulthood by steroid hormone manipulations. Based on this paper the notion was established that sex-typical reproductive behaviors are differentiated by brain changes induced by perinatal androgens or their metabolites. At that time Beach (reviewed in (Beach, 1971; Beach, 1981) challenged the notion that this sexual differentiation was dependent on the organization of the brain by steroid hormones but rather it was the perinatal effect of steroids on the development of effector organs such as the penis and other peripheral organs that was critical for the sexual differentiation of these critical reproductive behaviors. Our data show that like so many controversies in science there is evidence for both positions. It seems clear that the neural circuit regulating song is sexually differentiated at least in part by perinatal steroid hormone action but we establish here that a peripheral effector organ, such as the syrinx, also plays a critical role in explaining sexually differentiated courtship song.

## Online supporting information

**Fig. S1. Additional song features that were recorded and analyzed in male and female canaries that were treated with Silastic™ implants filled with testosterone (T) or with testosterone plus estradiol (T+E2) or left empty as control (C)**. Bar graphs represent the mean ± SEM of individual results that are the average of data collected during the 6 weeks of recording. Individual data points are also indicated. Data were analyzed by two-way ANOVA with treatment (Trt) and Sex of the subjects as independent factors and results are summarized in the insert for each panel that is shaded in gray when significant effects were detected. (***=p<0.001, **=p<0.01, *=p<0.05, ns= not significant). Significant effects of treatment were further analyzed by Tukey’s multiple comparison tests and their results are expressed as follows: a= p<0.05 versus C group.

**Fig. S2. Trill rate and trill features quantified by a MATLAB script in songs recorded from male and fe-male canaries that were treated with Silastic™ implants filled with testosterone (T) or with testosterone plus estradiol (T+E2) or left empty as control (C)**. The different graphs represent the mean ± SEM of individ-ual results collected during the 6 successive weeks of recording. Data were analyzed by two two-way ANOVAs separately for males and females with the treatment (TRT) and Time as factors and results, when significant, are summarized at the top left for each panel. (***=p<0.001, **=p<0.01, *=p<0.05). Significant effects of treatments were further analyzed by Tukey’s multiple comparison tests and their results are expressed as follows: aaa (or bbb)= p<0.001 versus C (or T) group, aa (or bb)= p<0.01 versus C (or T) group, a (or b)= p<0.05 versus C (or T) group. Significant interactions are described in the text.

***Fig. S3. Trade off between trill rate and trill bandwidth as observed separately in songs of all males (M; left panel) and all females (F; right panel) canaries that were treated with Silastic implants filled with testosterone (T) or with testosterone plus estradiol (T+E2)***.

***Fig. S4. Trade off between trill rate and trill bandwidth as observed separately in individual subjects that had produced at least 10 trills during the entire experiment***.

***Fig. S5. Differential effects in males and females of exogenous testosterone (T) associated or not with estradiol (E2) on the fiber diameter in the 4 quadrants of the syrinx (Q1 to Q4) (A) and on the overall fiber density (number/mm***^***2***^***) over the 4 quadrants (B)***. *Data were analyzed by two-way ANOVAs with treatment (Trt) and Sex of the subjects as independent factors and results are summarized in the insert for each panel. (***=p<0*.*001, **=p<0*.*01, *=p<0*.*05)*.

**Table S1. Trill analysis: Means of 6 weeks. The table presents the results (F with degrees of freedom and associated probabilities) of the two-way ANOVAs of the means of data collected over the six weeks of experiment**.

**Table S2. Trill analysis: group differences**. The table presents the results (F with degrees of freedom and associated probabilities) of the two-way ANOVAs of data collected each week to assess treatment effects separately in males and females.

**Table S3. Trill analysis: Sex differences**. The table presents the results (F with degrees of freedom and asso-ciated probabilities) of the two-way ANOVAs of data collected each week separately to assess sex differences separately in each experimental group.

## Acknowledgments

This work was supported by a Grant from the National Institute of Neurological Disorders and Stroke Grant RO1NS104008 (to G.F.B., J.B., and C.A.C.). We thank Pr. Robert Dooling and Ed. Smith, department of Psychology, University of Maryland in College Park for providing the MATLAB script used to analyze canary song. C.A.C. is F.R.S.-FNRS Research Director.

## Authors contributions

**Conceptualization:** Ednei B. dos Santos, Gregory F. Ball, Charlotte A. Cornil, Jacques

Balthazart

**Investigation:** Ednei B. dos Santos

**Formal analysis:** Ednei B. dos Santos, David M. Logue, Jacques Balthazart

**Writing-original draft:** Ednei B. dos Santos, Jacques Balthazart

**Writing-review & edition:** Ednei B. dos Santos, David M. Logue,GFB, Charlotte A. Cornil,

Jacques Balthazart

## References

Adam, I., Elemans, C.P.H., 2019. Vocal Motor Performance in Birdsong Requires Brain-Body Interaction. eNeuro 6.

Alvarez-Borda, B., Nottebohm, F., 2002. Gonads and singing play separate, additive roles in new neuron recruitment in adult canary brain. J Neurosci 22, 8684–8690.

Alward, B.A., Balthazart, J., Ball, G.F., 2013. Differential effects of global versus local testosterone on singing behavior and its underlying neural substrate. Proceedings of the National Academy of Sciences of the United States of America 110, 19573–19578.

Alward, B.A., Madison, F.N., Gravley, W.T., Ball, G.F., 2016. Antagonism of syringeal androgen receptors reduces the quality of female-preferred male song in canaries. Animal Behaviour 119, 201–212.

Andersson, M., 1994. Sexual selection. Princeton University Press, Princeton NJ.

Appeltants, D., Ball, G.F., Balthazart, J., 2003. Song activation by testosterone is associated with an increased catecholaminergic innervation of the song control system in female canaries. Neuroscience 121, 801–814.

Ball, G.F., Balthazart, J., 2007. The neuroendocrinology and neurochemistry of birdsong, in: Lajtha, A. (Ed.), Handbook of Neurochemistry and Molecualr Neurobiology, 3rd Edition, J.D. Blaustein (Volume Ed.). Springer, New York, pp. 419–457.

Beach, F.A., 1971. Hormonal factors controlling the differentiation, development and display of copulatory behavior in the ramstergig and related species, The Biopsychology of Development. Academic Press, New York,pp. 249–296.

Beach, F.A., 1981. Historical origins of modern research on hormones and behavior. Horm Behav 15, 325–376.

Cade, B.S., Noon, B.R., 2003. A gentle introduction to quantile regression for ecologists. Front. Ecol. Environ. 1, 412–420.

Catchpole, C.K., Slater, P.J.B., 2018. Bird song. Biological themes and Variations. Cambridge University Press, Cambridge UK.

Christensen, L.A., Allred, L.M., Goller, F., Meyers, R.A., 2017. Is sexual dimorphism in singing behavior related to syringeal muscle composition?. Auk 134, 710–720.

Collins, S., 2004. Vocal fighting and flirting: the functions of birdsong, in: Marler, P., Slabbekoorn, H. (Eds.), Nature’s Lusic, The Science of Birdsong. Elsevier, Amsterdam, pp. 39–79.

Cornez, G., Collignon, C., Muller, W., Cornil, C.A., Ball, G.F., Balthazart, J., 2020. Development of Perineuronal Nets during Ontogeny Correlates with Sensorimotor Vocal Learning in Canaries. eNeuro 7.

Dos Santos, E.B., Ball, G.F., Cornil, C.A., Balthazart, J., 2022. Treatment with androgens plus estrogens cannot reverse sex differences in song and the song control nuclei in adult canaries. Horm Behav 143, 105197.

Draganoiu, T.I., Nagle, L., Kreutzer, M., 2002. Directional female preference for an exaggerated male trait in canary (Serinus canaria) song. Proc Biol Sci 269, 2525–2531.

Elemans, C.P., Mead, A.F., Rome, L.C., Goller, F., 2008. Superfast vocal muscles control song production in songbirds. PLoS One 3, e2581.

Fuxjager, M.J., Eaton, J., Lindsay, W.R., Salwiczek, L.H., Rensel, M.A., Barske, J., Sorenson, L., Day, L.B., Schlinger, B.A., 2015. Evolutionary patterns of adaptive acrobatics and physical performance predict expression profiles of androgen receptor - but not oestrogen receptor -in the forelimb musculature. Funct Ecol 29, 1197–1208.

Geraci, M., 2014. Linear Quantile Mixed Models: The lqmm Package for Laplace Quantile Regression. J. Stat. Softw. 57, 1–29.

Goldman, S.A., Nottebohm, F., 1983. Neuronal production, migration, and differentiation in a vocal control nucleus of the adult female canary brain. Proceedings of the National Academy of Sciences of the United States of America 80, 2390–2394.

Goller, F., 2022. Vocal atheletics - from birdsong production mechanisms to sexy songs. Anim Behav 184, 173–184.

Griffiths, R., Double, M.C., Orr, K., Dawson, R.J., 1998. A DNA test to sex most birds. Mol Ecol 7, 1071–1075.

Haakenson, C.M., Madison, F.N., Ball, G.F., 2019. Effects of Song Experience and Song Quality on Immediate Early Gene Expression in Female Canaries (Serinus canaria). Dev Neurobiol 79, 521–535.

Hartog, T.E., Dittrich, F., Pieneman, A.W., Jansen, R.F., Frankl-Vilches, C., Lessmann, V., Lilliehook, C., Goldman, S.A., Gahr, M., 2009. Brain-derived neurotrophic factor signaling in the HVC is required for testosterone-induced song of female canaries. J Neurosci 29, 15511–15519.

Kirkpatrick, M., Ryan, M.J., 1991. The evolution of mating preferences and the paradox of the lek. NAture 350, 33–38.

Leboucher, G., Kreutzer, M., Dittami, J., 1994. Copulation-solicitation displays in female canaries (Serinus canaria): are oestradiol implants necessary? Ethology 97, 190–197.

Leitner, S., Voigt, C., Metzdorf, R., Catchpole, C.K., 2005. Immediate early gene (ZENK, Arc) expression in the auditory forebrain of female canaries varies in response to male song quality. Journal of neurobiology 64, 275–284.

Li, X.C., Jarvis, E.D., Alvarez-Borda, B., Lim, D.A., Nottebohm, F., 2000. A relationship between behavior, neurotrophin expression, and new neuron survival. PNAS 97, 8584–8589.

Logue, D.M., Sheppard, J.A., Walton, B., Brinkman, B.E., Medina, O.J., 2020. An analysis of avian vocal performance at the note and song levels. Bioacoustics 29, 709–730.

Louissaint, A.J., Rao, S., Leventhal, C., Goldman, S.A., 2002. Coordinated interaction of neurogenesis and angiogenesis in the adult songbird brain. Neuron 34, 945–960.

Madison, F.N., Rouse, M.L., Jr., Balthazart, J., Ball, G.F., 2015. Reversing song behavior phenotype: Testosterone driven induction of singing and measures of song quality in adult male and female canaries (Serinus canaria). Gen Comp Endocrinol 215, 61–75.

Maxwell, A., Adam, I., Larsen, P.S., Sorensen, P.G., Elemans, C.P.H., 2021. Syringeal vocal folds do not have a voice in zebra finch vocal development. Sci Rep 11, 6469.

Monbureau, M., Barker, J.M., Leboucher, G., Balthazart, J., 2015. Male song quality modulates c-Fos expression in the auditory forebrain of the female canary. Physiology & behavior 147, 7–15.

Nottebohm, F., Arnold, A.P., 1976. Sexual dimorphism in vocal control areas of the songbird brain. Science 194, 211–213.

Owren, M.J., Berkowitz, M., Bachorowski, J.A., 2007. Listeners judge talker sex more efficiently from male than from female vowels. Percept Psychophys 69, 930–941.

Phoenix, C.H., Goy, R.W., Gerall, A.A., Young, W.C., 1959. Organizational action of prenatally administered testosterone propionate on the tissues mediating behavior in the female guinea pig. Endocrinology 65, 369–382.

Podos, J., 1997. A Performance Constraint on the Evolution of Trilled Vocalizations in a Songbird Family (Passeriformes: Emberizidae). Evolution 51, 537–551.

Podos, J., 2001. Correlated evolution of morphology and vocal signal structure in Darwin’s finches. Nature 409, 185–188.

Podos, J., Lahti, D.C., Moseley, D.L., 2009. Vocal Performance and Sensorimotor Learning in Songbirds. Advances in the Study of Animal Behavior (Academic Press),159–195.

Podos, J., Sung, H.-C., 2020. Vocal Performance in Songbirds: From Mechanisms to Evolution, in: Sakata, J.T., Woolley, S.C., Fay, R.R., Popper, A.N. (Eds.), The Neuroethology of Birdsong, Springer Handbook of Auditory Research., pp. 245–268.

Prince, B., Riede, T., Goller, F., 2011. Sexual dimorphism and bilateral asymmetry of syrinx and vocal tract in the European starling (Sturnus vulgaris). J Morphol 272, 1527–1536.

Rehsteiner, U., Geisser, H., Reyer, H., 1998. Singing and mating success in water pipits: one specific song element makes all the difference. Anim Behav 55, 1471–1481.

Sakata, J.T., Vehrencamp, S.L., 2012. Integrating perspectives on vocal performance and consistency. J Exp Biol 215, 201–209.

Sartor, J.J., Balthazart, J., Ball, G.F., 2005. Coordinated and dissociated effects of testosterone on singing behavior and song control nuclei in canaries (Serinus canaria). Hormones and Behavior 47, 467–476.

Schindelin, J., Arganda-Carreras, I., Frise, E., Kaynig, V., Longair, M., Pietzsch, T., Preibisch, S., Rueden, C., Saalfeld, S., Schmid, B., Tinevez, J.Y., White, D.J., Hartenstein, V., Eliceiri, K., Tomancak, P., Cardona, A., 2012. Fiji: an open-source platform for biological-image analysis. Nat Methods 9, 676–682.

Schlinger, B.A., Brenowitz, E.A., 2017. Neural and hormonal control of birdsong, in: Pfaff, D.W., Joels, M. (Eds.), Hormones, brain and behavior, 3rd ed. Academic Press, Oxford, pp. 255–290.

Shevchouk, O.T., Ball, G.F., Cornil, C.A., Balthazart, J., 2017. Studies of HVC Plasticity in Adult Canaries Reveal Social Effects and Sex Differences as Well as Limitations of Multiple Markers Available to Assess Adult Neurogenesis. PLoS One 12, e0170938.

Suthers, R.A., Vallet, E., Kreutzer, M., 2012. Bilateral coordination and the motor basis of female preference for sexual signals in canary song. J Exp Biol 215, 2950–2959.

Team, R., 2021. RStudio: integrated development for R.

Tobet, S.A., Fox, T.O., 1992. Sex differences in neuronal morphology influenced hormonally throughout life. Handbook of behavioral neurobiology. Vol 11. Sexual differentiation, in: Gerall, A.A., Moltz, H., Ward, I.L. (Eds.), 0 ed. Plenum Press, New York, pp. 41–83.

Tramontin, A.D., Wingfield, J.C., Brenowitz, E.A., 2003. Androgens and estrogens induce seasonal-like growth of song nuclei in the adult songbird brain. Journal of neurobiology 57, 130–140.

Vallet, E., Beme, I.I., Kreutzer, M., 1998. Two-note syllables in canary songs elicit high levels of sexual display. Anim Behav 55, 291–297.

Vallet, E., Kreutzer, M., 1995. Female canaries are sexually responsive to special song phrases. Anim.Behav. 49, 1603–1610.

Veney, S.L., Wade, J., 2004. Steroid receptors in the adult zebra finch syrinx: a sex difference in androgen receptor mRNA, minimal expression of estrogen receptor alpha and aromatase. Gen Comp Endocrinol 136, 192–199.

Veney, S.L., Wade, J., 2005. Post-hatching syrinx development in the zebra finch: an analysis of androgen receptor, aromatase, estrogen receptor alpha and estrogen receptor beta mRNAs. J Comp Physiol A Neuroethol Sens Neural Behav Physiol 191, 97–104.

Wade, J., Buhlman, L., 2000. Lateralization and effects of adult androgen in a sexually dimorphic neuromuscular system controlling song in zebra finches. J.Comp.Neurol. 426, 154–164.

Wickham, H., Chang, W., 2008. ggplot2: an implementation of the grammar of graphics (https://ggplot2-book.org/introduction.html).

Wilson, D.R., Bitton, P.-P., Podos, J., 1997. Uneven Sampling and the Analysis of Vocal Performance Constraints. Am. Nat. 183, 214–228.

Yamamura, T., Barker, J.M., Balthazart, J., Ball, G.F., 2011. Androgens and estrogens synergistically regulate the expression of doublecortin and enhance neuronal recruitment in the song system of adult female canaries. Journal of Neuroscience 31, 9649–9657.

