## Supporting figures and tables for "Does the syrinx, a peripheral structure, constrain effects of sex steroids on behavioral sex reversal in adult canaries?"

**This PDF file includes:**

Supplementary results
Figures S1 to S5
Tables S1 to S3

**Trill features quantified in the average data collected over the 6 weeks of experiment**

In addition to the 3 dependent variables discussed in the main text, the MATLAB routine quantified 9 additional characteristics of the trills. Averages of these data over the six weeks of experiments are summarized in figure S1 and results of the analysis by two-way ANOVA of these data is presented in table S1.

As mentioned in the main text, three of these additional variables were significantly affected by the treatments, namely the number of segments per trill, the segment duration and the number of fast trills with more than 17 segments per seconds but other variable were not. In addition, none of these variables were associated with a sex difference nor with an interaction between treatment and sex.

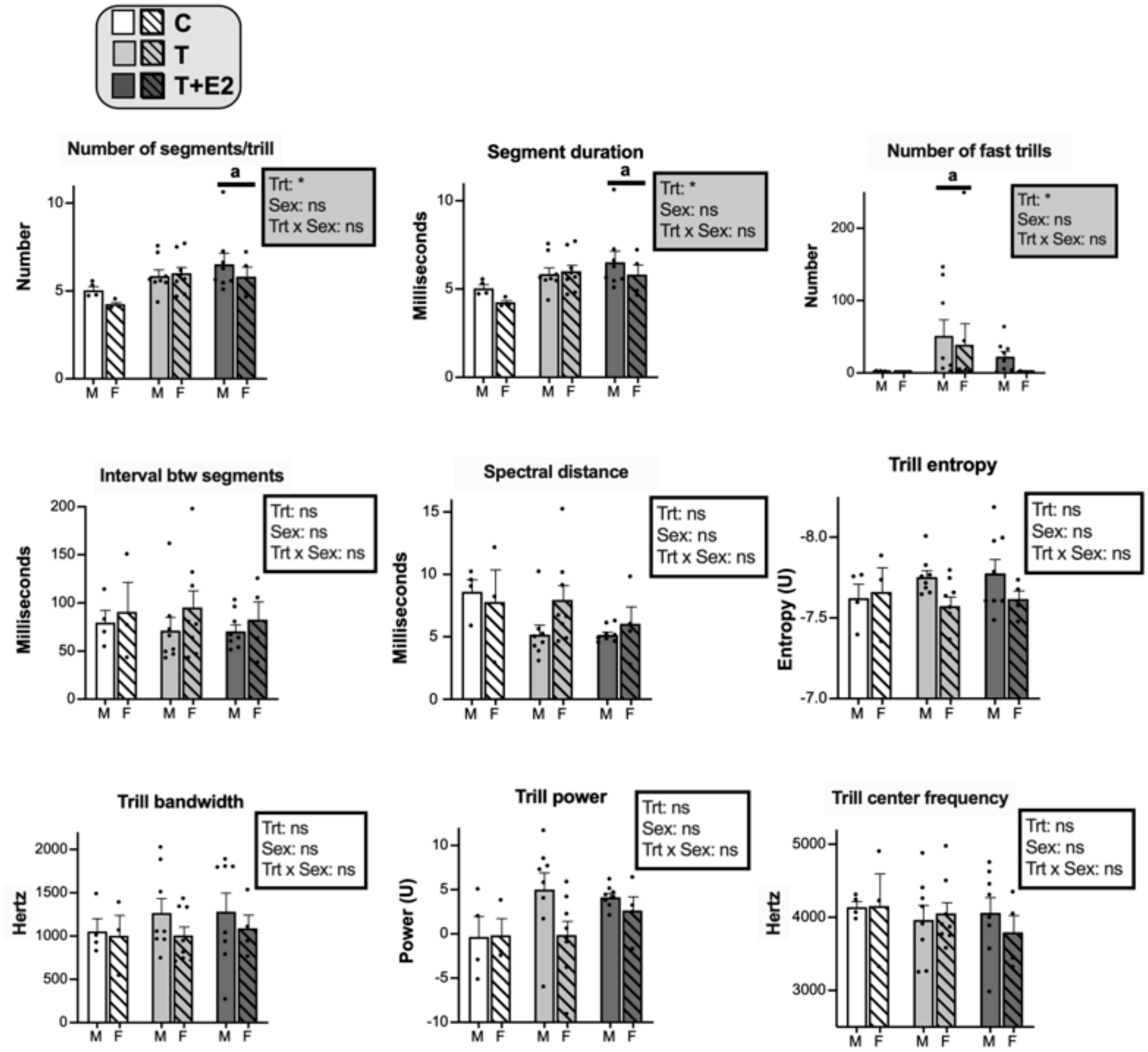


**Fig. S1.** Additional song features that were recorded and analyzed in male and female canaries that were treated with Silastic™ implants filled with testosterone (T) or with testosterone plus estradiol (T+E2) or left empty as control (C). Bar graphs represent the mean ± SEM of individual results that are the average of data collected during the 6 weeks of recording. Individual data points are also indicated. Data were analyzed by two-way ANOVA with treatment (Trt) and Sex of the subjects as independent factors and results are summarized in the insert for each panel that is shaded in gray when significant effects were detected. (***=p<0.001, **=p<0.01, *=p<0.05, ns= not significant). Significant effects of treatment were further analyzed by Tukey's multiple comparison tests and their results are expressed as follows: a= p<0.05 versus C group.

**Table S1.** Trill analysis: Means of 6 weeks. The table presents the results (F with degrees of freedom and associated probabilities) of the two-way ANOVAs of the means of data collected over the six weeks of experiment.

| | **Treatment** | | | **Sex** | | | **Interaction** | | |
|---|---|---|---|---|---|---|---|---|---|
| **Variable** | $\mathbf{F}_{df}$ | **p** | $\eta_p^2$ | $\mathbf{F}_{df}$ | **p** | $\eta_p^2$ | $\mathbf{F}_{df}$ | **p** | $\eta_p^2$ |
| **Number of trills** | $F_{2,42}$ = 13.900 | **<0.001** | 0.40 | $F_{1,42}$ = 8.815 | **0.005** | 0.17 | $F_{2,42}$ = 1.567 | 0.221 | 0.07 |
| **% Time trilling** | $F_{2,42}$ = 8.288 | **<0.001** | 0.28 | $F_{1,42}$ = 9.267 | **0.004** | 0.18 | $F_{2,42}$ = 1.656 | 0.203 | 0.07 |
| **Trill duration** | $F_{2,42}$ = 20.820 | **<0.001** | 0.50 | $F_{1,42}$ =18.66 | **<0.001** | 0.31 | $F_{2,42}$ = 4.337 | **0.019** | 0.17 |
| **Nbr Segments/trill** | $F_{2,29}$ = 3.558 | **0.041** | 0.20 | $F_{1,29}$ = 1.261 | 0.271 | 0.04 | $F_{2,29}$ = 0.575 | 0.569 | 0.04 |
| **Segment duration** | $F_{2,29}$ = 3.558 | **0.041** | 0.20 | $F_{1,29}$ = 1.261 | 0.271 | 0.04 | $F_{2,29}$ = 0.575 | 0.569 | 0.04 |
| **Fast trills (>17 segm.. $S^{-1}$)** | $F_{2,42}$ = 4.259 | **0.021*** | 0.17 | $F_{1,42}$ = 0.894 | 0.350 | 0.02 | $F_{2,42}$ = 0.240 | 0.788 | 0.01 |
| **Interval btw trills** | $F_{2,29}$ = 0.134 | 0.875 | 0.01 | $F_{1,29}$ = 1.022 | 0.320 | 0.03 | $F_{2,29}$ = 0.100 | 0.905 | 0.01 |
| **Interval btw segments** | $F_{2,29}$ = 2.036 | 0.149 | 0.12 | $F_{1,29}$ = 0.799 | 0.379 | 0.03 | $F_{2,29}$ = 1.199 | 0.316 | 0.08 |
| **Trill entropy** | $F_{2,29}$ = 0.196 | 0.823 | 0.01 | $F_{1,29}$ = 2.402 | 0.132 | 0.08 | $F_{2,29}$ = 0.849 | 0.43 | 0.06 |
| **Trill bandwidth** | $F_{2,29}$ = 0.261 | 0.772 | 0.02 | $F_{1,29}$ =1.183 | 0.286 | 0.04 | $F_{2,29}$ = 0.131 | 0.877 | 0.01 |
| **Trill power** | $F_{2,29}$ = 1.694 | 0.201 | 0.10 | $F_{1,29}$ = 2.285 | 0.141 | 0.07 | $F_{2,29}$ = 1.223 | 0.309 | 0.08 |
| **Tril center frequency** | $F_{2,29}$ = 0.352 | 0.706 | 0.02 | $F_{1,29}$ = 0.125 | 0.726 | 0.00 | $F_{2,29}$ = 0.367 | 0.696 | 0.02 |

### Effects of time

Trills were rarely produced by females and castrated males and they only appeared progressively after birds had been treated for some time with exogenous steroids. As a consequence, these vocalizations were frequently absent in song recordings and this prevented in multiple cases the statistical analysis of their characteristics and modifications as a function of time. Average data are presented in **Fig. S2.**

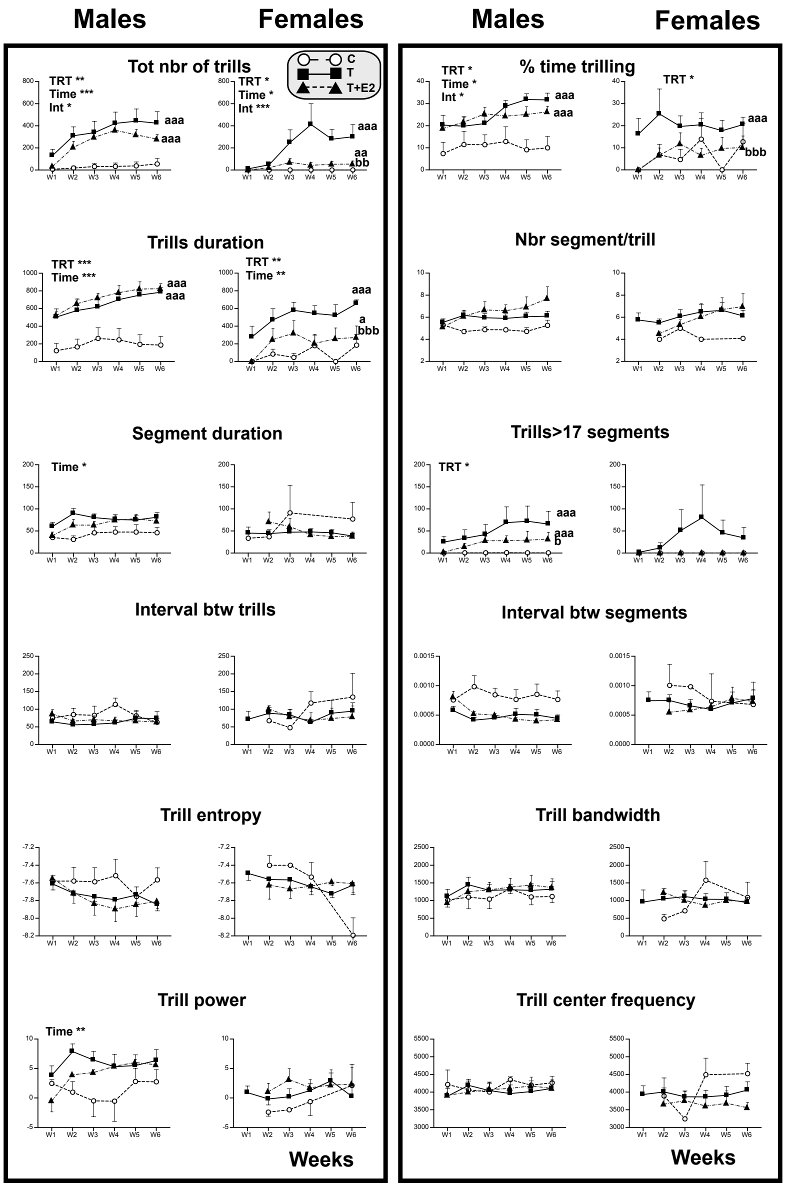


**Fig. S2.** Trill rate and trill features quantified by a MATLAB script in songs recorded from male and female canaries that were treated with Silastic™ implants filled with testosterone (T) or with testosterone plus estradiol (T+E2) or left empty as control (C). The different graphs represent the mean ± SEM of individual results collected during the 6 successive weeks of recording. Data were analyzed by two two-way ANOVAs separately for males and females with the treatment (TRT) and Time as factors and results, when significant, are summarized at the top left for each panel. (***=$p<0.001$, **=$p<0.01$, *=$p<0.05$). Significant effects of treatments were further analyzed by Tukey's multiple comparison tests and their results are expressed as follows: aaa (or bbb)= $p<0.001$ versus C (or T) group, aa (or bb)= $p<0.01$ versus C (or T) group, a (or b)= $p<0.05$ versus C (or T) group. Significant interactions are described in the text.

**Within sexes.** Results of the statistical analysis of the effects of the endocrine treatments as a function of time, when sufficient number of data were available, are summarized for each sex separately in **Table S2** (M=males, F=females). Most of these effects developed progressively and were thus associated with a significant effect of time; the percentage time trilling in females and numbers of fast trills in males were the only exceptions (see **Table S2** for detail). A few significant interactions between time and treatments were also detected (total number of trills in males and females, percentage time trilling and spectral distance in males only) and reflected the fact that steroids progressively affected trill characteristics that did not change in control birds.

**Table S2. Trill analysis: group differences.** The table presents the results (F with degrees of freedom and associated probabilities) of the two-way ANOVAs of data collected each week to assess treatment effects separately in males and females.

| **Variables** | | **Time** | | **Treatment** | | **Time x Treatment** | |
|---|---|---|---|---|---|---|---|
| | | $F_{df}$ | *P* | $F_{df}$ | *P* | $F_{df}$ | *P* |
| Number of trills | M | $F_{3.180, 66.79} = 12.77$ | **<0.0001*** ** | $F_{2, 21} = 9.324$ | **0.0013** ** | $F_{10, 105} = 2.488$ | **0.0102*** |
| | F | $F_{1.505, 31.01} = 4.876$ | **0.0219*** | $F_{2, 21} = 5.760$ | **0.0101*** | $F_{10, 103} = 3.408$ | **0.0007*** ** |
| % Time trilling | M | $F_{3.220, 66.97} = 3.530$ | **0.0172*** | $F_{2, 21} = 5.741$ | **0.0103*** | $F_{10, 104} = 1.915$ | **0.0511*** |
| | F | $F_{2.761, 56.87} = 1.611$ | 0.2 | $F_{2, 21} = 4.316$ | **0.0269*** | $F_{10, 103} = 0.6868$ | 0.7346 |
| Trill duration | M | $F_{2.848, 59.81} = 7.485$ | **0.0003*** ** | $F_{2, 21} = 15.74$ | **<0.0001*** ** | $F_{10, 105} = 1.127$ | 0.3495 |
| | F | $F_{3.303, 68.04} = 5.320$ | **0.0017** ** | $F_{2, 21} = 9.204$ | **0.0013** ** | $F_{10, 103} = 0.8578$ | 0.5748 |
| Number of segments | M | $F_{1.667, 25.00} = 2.665$ | 0.0974 | $F_{2, 17} = 1.504$ | 0.2503 | $F_{10, 75} = 1.601$ | 0.1227 |
| Segment duration | M | $F_{2.146, 32.19} = 3.895$ | **0.028*** | $F_{2, 17} = 2.718$ | 0.0946 | $F_{10, 75} = 0.7492$ | 0.6762 |
| Fast trills (> 17 segm. $s^{-1}$) | M | $F_{2.482, 52.11} = 2.457$ | 0.0839 | $F_{2, 21} = 3.520$ | **0.0481*** | $F_{10, 105} = 0.8651$ | 0.5681 |
| | F | $F_{1.113, 22.92} = 1.076$ | 0.3184 | $F_{2, 21} = 1.493$ | 0.2476 | $F_{10, 103} = 1.067$ | 0.3945 |
| Interval duration | M | $F_{2.119, 31.78} = 0.6067$ | 0.5605 | $F_{2, 17} = 0.1220$ | 0.8859 | $F_{10, 75} = 1.596$ | 0.1243 |
| Spectral distance | M | $F_{2.255, 33.83} = 3.910$ | **0.0255*** | $F_{2, 17} = 6.104$ | **0.01*** | $F_{10, 75} = 2.312$ | **0.0198*** |
| Trill entropy | M | $F_{2.504, 37.55} = 1.560$ | 0.2197 | $F_{2, 17} = 0.9791$ | 0.3959 | $F_{10, 75} = 0.5284$ | 0.8649 |
| Trill bandwidth | M | $F_{2.804, 42.06} = 2.638$ | 0.0655 | $F_{2, 17} = 0.2654$ | 0.77 | $F_{10, 75} = 0.4826$ | 0.8963 |
| Trill power | M | $F_{2.037, 30.55} = 7.886$ | **0.0016** ** | $F_{2, 17} = 2.107$ | 0.1522 | $F_{10, 75} = 1.583$ | 0.1283 |
| Trill center frequency | M | $F_{3.612, 54.18} = 1.776$ | 0.1529 | $F_{2, 17} = 0.2229$ | 0.8025 | $F_{10, 75} = 0.4063$ | 0.9396 |

**Within treatment.** The frequent absence of trills in females and in C males often prevented the statistical analysis of the sex differences in trill characteristics over time. This was in fact possible only for dependent variables for which the absence of trills was meaningful and encoded as a zero in the data tables. For other dependent variables that could not be studied if trills were not produced (e.g., their entropy or bandwidth), sufficient data were only available in T treated birds; the T+E2 bird females produced much fewer trills than the T females. A few effects of time were also detected and concerned exclusively the T or T+E2 but since they mostly overlap with the effects of treatments detected in the average data, they will not be further discussed (see **Table S3** for detail). A single interaction between time and sex was finally detected and concerned the total number of trill in T+E2 birds: in this endocrine condition, trill numbers increased markedly with time in males but not in females.

**Table S3. Trill analysis: Sex differences.** The table presents the results (F with degrees of freedom and associated probabilities) of the two-way ANOVAs of data collected each week separately to assess sex differences separately in each experimental group.

| **Variables** | | **Time** | | **Sex** | | **Time x Sex** | |
|---|---|---|---|---|---|---|---|
| | | $F_{df}$ | *P* | $F_{df}$ | *P* | $F_{df}$ | *P* |
| Number of trills | C | $F_{1.003,\ 14.04} = 1.148$ | 0.3022 | $F_{1,\ 14} = 1.120$ | 0.3079 | $F_{5,\ 70} = 1.087$ | 0.3751 |
| | T | $F_{1.799,\ 25.18} = 8.152$ | **0.0024**** | $F_{1,\ 14} = 1.224$ | 0.2873 | $F_{5,\ 70} = 0.8238$ | 0.5369 |
| | T+E2 | $F_{3.126,\ 42.52} = 8.398$ | **0.0001***** | $F_{1,\ 14} = 40.87$ | **<0.0001***** | $F_{5,\ 68} = 4.005$ | **0.003**** |
| % Time trilling | C | $F_{1.877,\ 25.91} = 1.525$ | 0.2367 | $F_{1,\ 14} = 0.5322$ | 0.4777 | $F_{5,\ 69} = 0.8271$ | 0.5347 |
| | T | $F_{1.781,\ 24.93} = 0.9032$ | 0.4075 | $F_{1,\ 14} = 1.391$ | 0.2579 | $F_{5,\ 70} = 1.297$ | 0.275 |
| | T+E2 | $F_{2.670,\ 36.32} = 5.222$ | **0.0056**** | $F_{1,\ 14} = 16.72$ | **0.0011**** | $F_{5,\ 68} = 0.4246$ | 0.83 |
| Trill duration | C | $F_{1.917,\ 26.84} = 2.055$ | 0.1493 | $F_{1,\ 14} = 1.106$ | 0.3107 | $F_{5,\ 70} = 1.060$ | 0.3899 |
| | T | $F_{2.831,\ 39.64} = 4.899$ | **0.0062**** | $F_{1,\ 14} = 2.314$ | 0.1505 | $F_{5,\ 70} = 0.5183$ | 0.7616 |
| | T+E2 | $F_{2.277,\ 30.97} = 6.372$ | **0.0035**** | $F_{1,\ 14} = 24.99$ | **0.0002***** | $F_{5,\ 68} = 0.6634$ | 0.6524 |
| Number of segments | T | $F_{1.844,\ 20.65} = 2.240$ | 0.1347 | $F_{1,\ 14} = 0.02740$ | 0.8709 | $F_{5,\ 56} = 0.7092$ | 0.619 |
| Segment duration | T | $F_{1.661,\ 18.60} = 1.453$ | 0.2568 | $F_{1,\ 14} = 8.503$ | **0.0113*** | $F_{5,\ 56} = 1.279$ | 0.286 |
| Fast trills (> 17 segm. $s^{-1}$) | C | $F_{1.627,\ 22.77} = 0.4889$ | 0.5814 | $F_{1,\ 14} = 3.853$ | 0.0698 | $F_{5,\ 70} = 0.4889$ | 0.7834 |
| | T | $F_{1.498,\ 20.97} = 2.040$ | 0.163 | $F_{1,\ 14} = 0.1319$ | 0.7219 | $F_{5,\ 70} = 0.3549$ | 0.8774 |

| | | | | | | | |
|---|---|---|---|---|---|---|---|
| | T+E2 | $F_{1.654, 22.50} = 1.152$ | 0.3248 | $F_{1, 14} = 8.605$ | **0.0109*** | $F_{5, 68} = 1.117$ | 0.3597 |
| Interval duration | T | $F_{1.493, 16.72} = 0.7063$ | 0.4678 | $F_{1, 14} = 0.8150$ | 0.3819 | $F_{5, 56} = 0.2341$ | 0.9459 |
| Spectral distance | T | $F_{2.065, 23.13} = 0.7412$ | 0.4916 | $F_{1, 14} = 3.688$ | 0.0754 | $F_{5, 56} = 0.7099$ | 0.6185 |
| Trill entropy | T | $F_{3.206, 35.91} = 1.318$ | 0.2834 | $F_{1, 14} = 5.392$ | **0.0358*** | $F_{5, 56} = 0.3152$ | 0.9018 |
| Trill bandwidth | T | $F_{1.512, 16.93} = 0.6725$ | 0.4834 | $F_{1, 14} = 1.668$ | 0.2175 | $F_{5, 56} = 0.2041$ | 0.9595 |
| Trill power | T | $F_{1.834, 20.54} = 6.861$ | **0.0062**** | $F_{1, 14} = 4.111$ | 0.0621 | $F_{5, 56} = 2.346$ | 0.0528 |
| Trill center frequency | T | $F_{1.741, 19.50} = 0.6179$ | 0.5278 | $F_{1, 14} = 0.02645$ | 0.8731 | $F_{5, 56} = 0.2674$ | 0.929 |

**Additional information on the trade-off between trill rates (TR) and trill bandwidth (TBW).**
Separate plots of the relationships between TR and TBW in males and females separating in addition the T and T+E2 birds clearly indicated that different clusters of data points were present in the 4 groups of subjects further suggesting that these clusters might represent different types of syllables produced by different subjects (Fig S3).

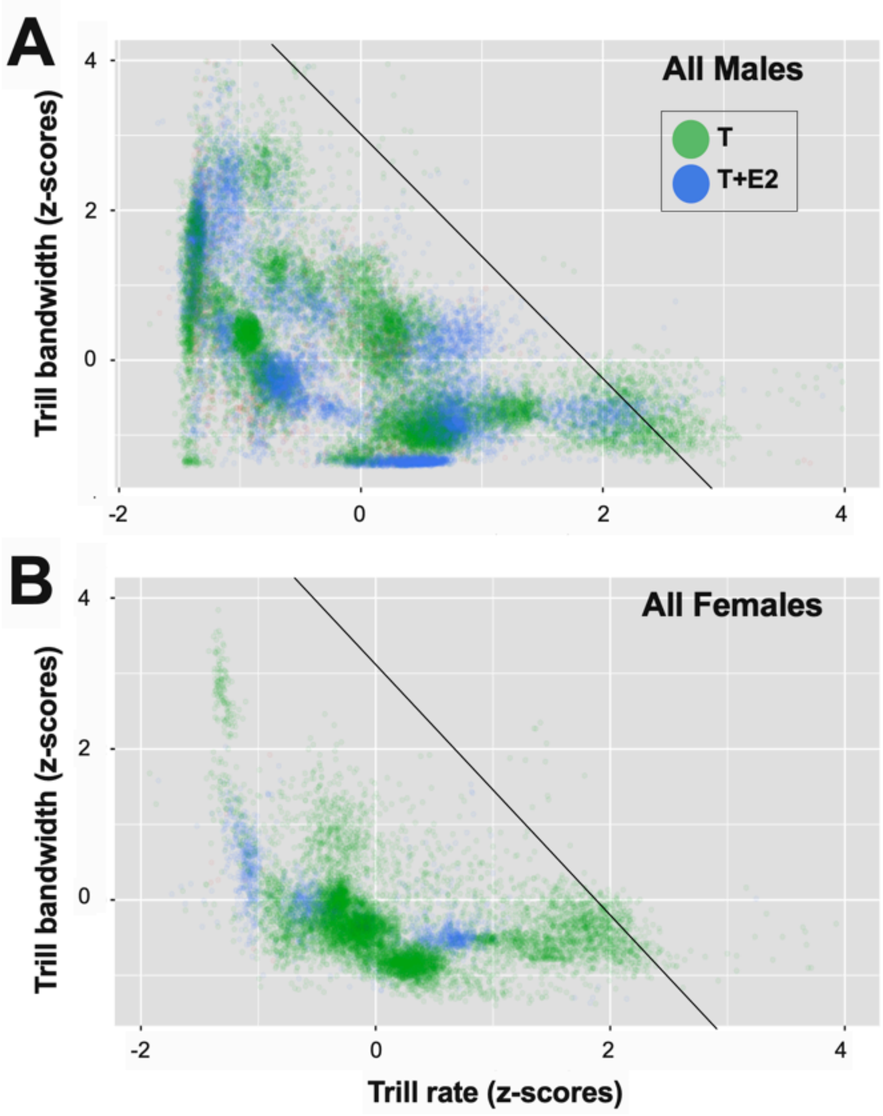


***Fig. S3.*** *Trade off between trill rate and trill bandwidth as observed separately in songs of all males (M; left panel) and all females (F; right panel) canaries that were treated with Silastic implants filled with testosterone (T) or with testosterone plus estradiol (T+E2).*

This suggestion was further reinforced by the separate plots of the trade-off in all individual subjects that had produced at least 10 trills during the entire experiment (Fig. S4). It can easily be observed that some clusters were present in multiple subjects whereas other clusters were specific to individual subjects.

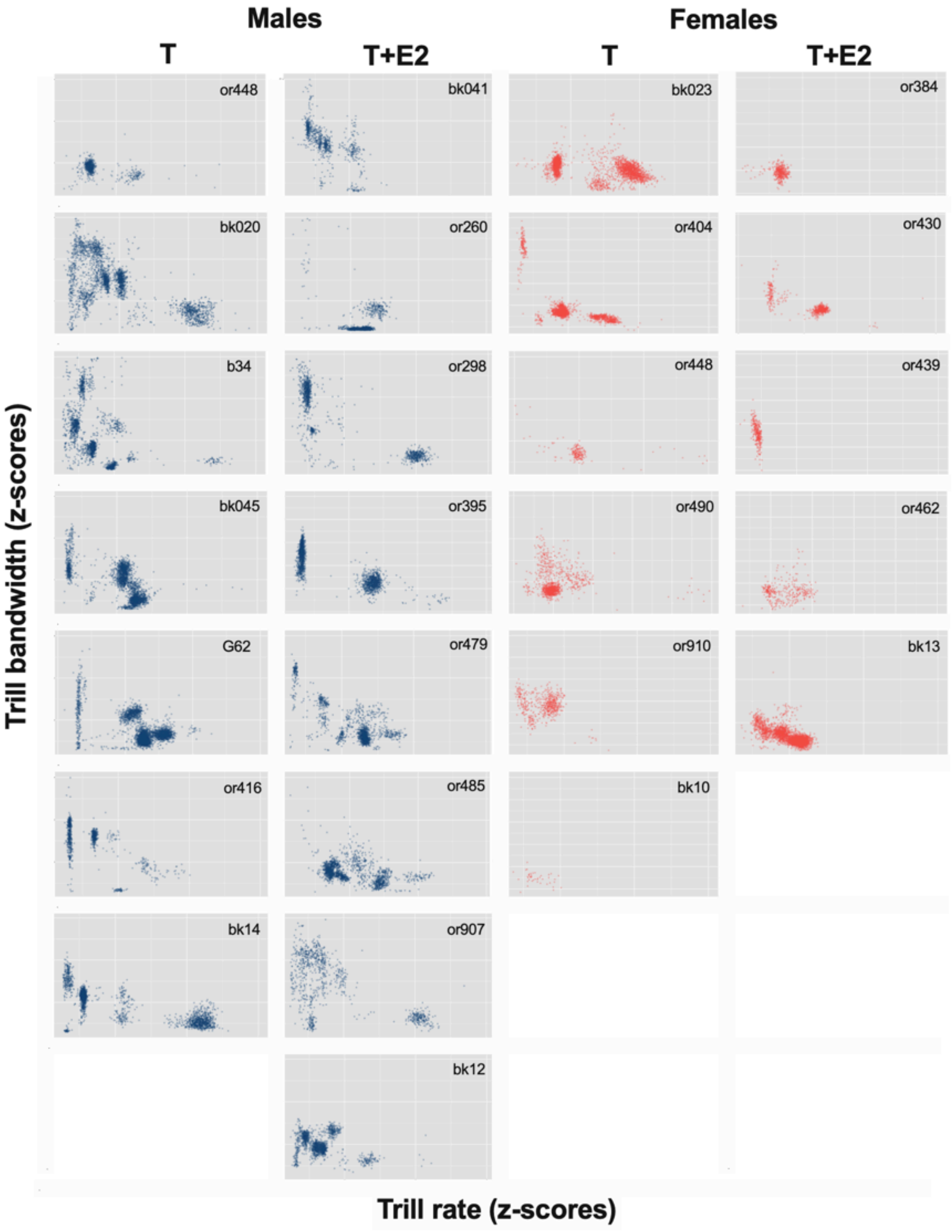


***Fig. S4.*** *Trade off between trill rate and trill bandwidth as observed separately in individual subjects that had produced at least 10 trills during the entire experiment.*

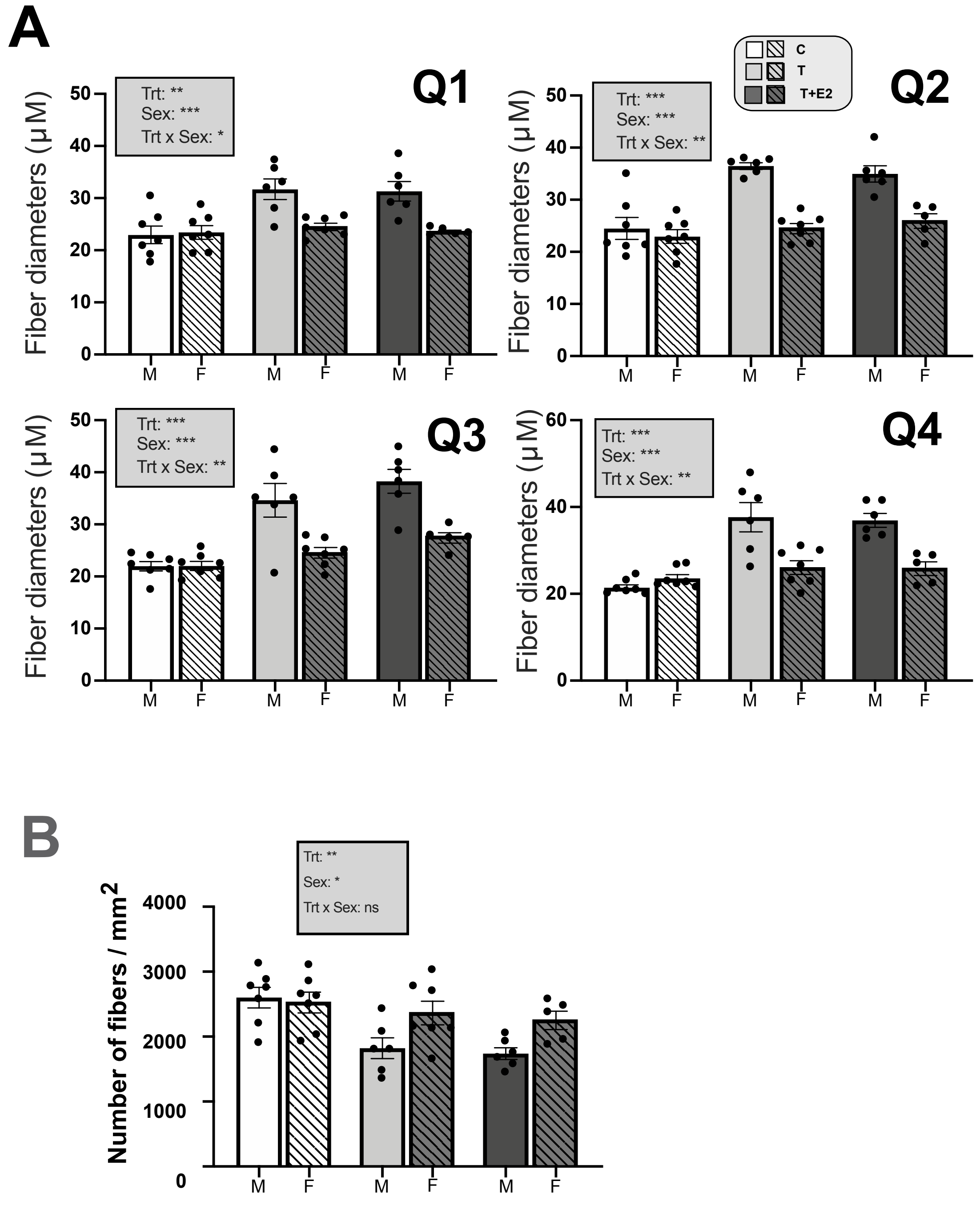


*Figure S5. Differential effects in males and females of exogenous testosterone (T) associated or not with estradiol (E2) on the fiber diameter in the 4 quadrants of the syrinx (Q1 to Q4) (A) and on the overall fiber density (number/mm$^2$) over the 4 quadrants (B). Data were analyzed by two-way ANOVAs with treatment (Trt) and Sex of the subjects as independent factors and results are summarized in the insert for each panel. (***=p<0.001, **=p<0.01, *=p<0.05).*
